# Ploidy modulates cell size and metabolic rate in *Xenopus* embryos

**DOI:** 10.1101/2022.10.17.512616

**Authors:** Clotilde Cadart, Julianne Bartz, Gillian Oaks, Martin Liu, Rebecca Heald

**Affiliations:** Department of Molecular and Cell Biology; Berkeley, CA 94720, USA

## Abstract

A positive correlation between genome size and cell size is well documented, but impacts on animal physiology are poorly understood. In *Xenopus* frogs, the number of genome copies (ploidy) varies across species and can be manipulated within a species. Here we show that triploid tadpoles contain fewer, larger cells than diploids and consume oxygen at a lower rate. Treatments that altered cell membrane stability or electrical potential abolished this difference, suggesting that a decrease in total cell surface area reduces basal energy consumption in triploids. Comparison of *Xenopus* species that evolved through polyploidization revealed that metabolic differences emerged during development when cell size scaled with genome size. Thus, ploidy affects metabolism by altering the cell surface area to volume ratio in a multicellular organism.

**One-Sentence Summary:** The amount of DNA per cell in a vertebrate modulates basal metabolism by altering cell size and plasma membrane energetics.

## Main Text

An increase in genome copy number or ploidy is a central phenomenon in biology that occurs during the development of specific tissues (*1*), is a first step in tumorigenesis (*2*) and is a major driving force of speciation throughout evolution (*3*). In these various contexts, the benefits of polyploidization are diverse, but one fundamental question is how ploidy impacts energy expenditure of the tissue, tumor, or organism. While the consequences of polyploidy on plant fitness have been well documented (*4*), they remain poorly characterized in animals. A potential mechanism of ploidy effects is through changes in cell size, a ubiquitous consequence of polyploidization in yeast (*5*, *6*) and vertebrate muscle cells (*7*). At the single-cell level, research on experimentally induced polyploid yeast (*6*) or animal cells (*8*), as well as enlarged animal cells (*9*–*11*) is beginning to unravel unexpected connections among ploidy, cell size, fitness, and metabolism (*12*) but has not yet revealed consequences for tissues. For example, while numerous studies have investigated metabolic shifts and allometric relationships in cancer (*13*), how changes in cell size and ploidy, both hallmarks of tumorigenesis (*14*), alter tumor metabolism remains poorly understood. Importantly, species comparisons have demonstrated a negative association between genome size and developmental rate in amphibians (*15*), as well as a more debated negative correlation with metabolic rate in adult birds and amphibians (*16*, *17*), suggesting that genome size is connected to whole-organism physiology. Despite the fundamental importance of understanding how genome copy number impacts animal physiology, there is no established system for quantifying the metabolic effects of varying ploidy *in vivo* to reveal underlying mechanisms and consequences.

To investigate the impact of ploidy on metabolism, we focused on frog embryos of the genus *Xenopus*, since a connection with developmental rate across Amphibian species has already been reported (*15*) and whole-genome duplications frequently occur in nature (*18*, *19*). We manipulated fertilization conditions to produce triploid *X. laevis* embryos and compared them to diploid embryos from the same clutch of eggs (Fig. 1, A-C). Triploid embryos accumulated mass similarly to diploids except for a transient ~5% decrease at day 3 post fertilization (p.f.) (Fig. 1D, Table S1) and had comparable developmental and survival rates (Fig. S1A-B). To measure cell size, we stained for cortical actin (Fig. 1E) and quantified the area of multi-ciliated cells present on the surface of stage 37 and 41 embryos. Triploids displayed an increase in cell area reflecting a linear relationship between spherical cell volume and ploidy (Fig. 1F). Therefore, triploid *X. laevis* embryos provide a means to evaluate the effects of a 50% genome size and comparable cell size increase on metabolism in a normally developing vertebrate.

**Fig. 1.**
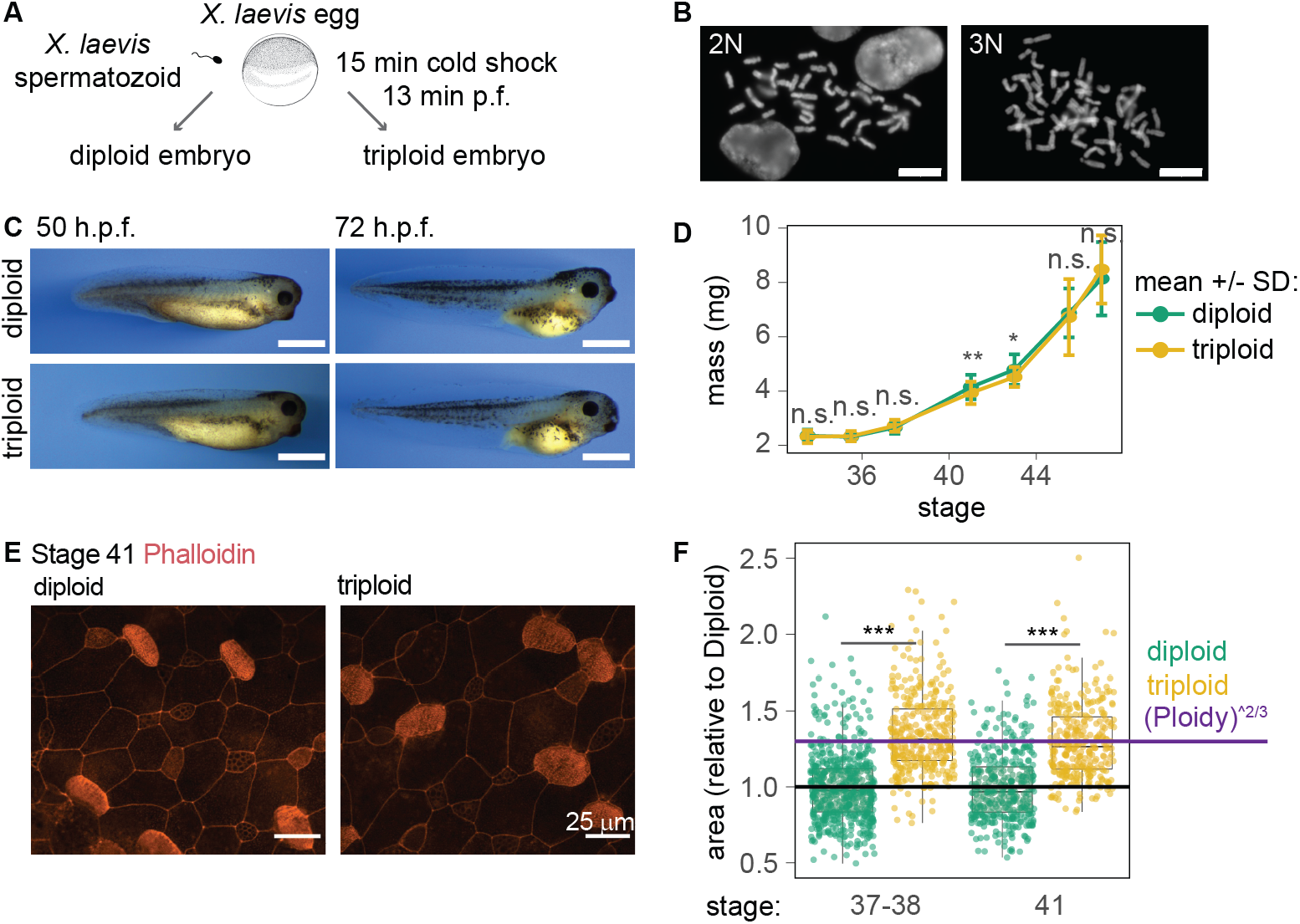
Triploid *X. laevis* develop normally. (**A**) Triploid *X. laevis* were obtained by blocking polar body extrusion post fertilization (p.f.) using a cold shock. (**B**) The 1.5-fold increase in chromosome number in triploid (3N) compared to diploid (2N) embryos was verified by metaphase chromosome spreads. Scale bar = 10 μm. (**C**) Representative images of diploid and triploid embryos obtained from the same clutch of eggs. Scale bar = 1 mm. Compared to diploids, triploids showed a comparable mass increase except for a transient ~ 5% reduction during day 3 (**D**) (19 clutches, n=10 to 82 embryos per ploidy and stage, details in Table S1). (**E**) TMR-phalloidin staining of actin in ventral epithelial cells of stage 41 diploid and triploid *X. laevis* embryos showing cell outlines and strong actin enrichment at the surface of multiciliated cells. (**F**) Area of diploid and triploid multiciliated cells in embryos at stage 37-38 and 41, normalized to diploids of each clutch. (3 clutches, n>300 for each condition). **(D), (F)**, Welch two sample t-test comparing the means, *: p<0.5, **: p<0.01,***: p<0.001.

Metabolic rate *B* and body mass *m* are related across all animals by the famous Kleiber’s law (*20*): *B* = *B*_0_*m^α^* with an allometric exponent α value that is typically between 0.6 and 0.75 for adult animals (*21*, *22*), but has not been characterized during development. Using a non-invasive detection system, we measured oxygen consumption rate (OCR), the most commonly used proxy for whole-organism metabolism (*23*), of single embryos in glass vials filled with water by recording over 5-7 hours (Fig. 2A, Fig. S2A-H). OCR measurement of over 250 diploid embryos from day 2 to day 5 post fertilization showed that metabolic rate increased sub-linearly with body mass, with a scaling exponent (*α* = 0.58 ± 0.02) consistent with Kleiber’s law (Fig. 2B, SI and Table S2). Thus, as in adults, allometric metabolic scaling occurs in early vertebrate development.

**Fig. 2:**
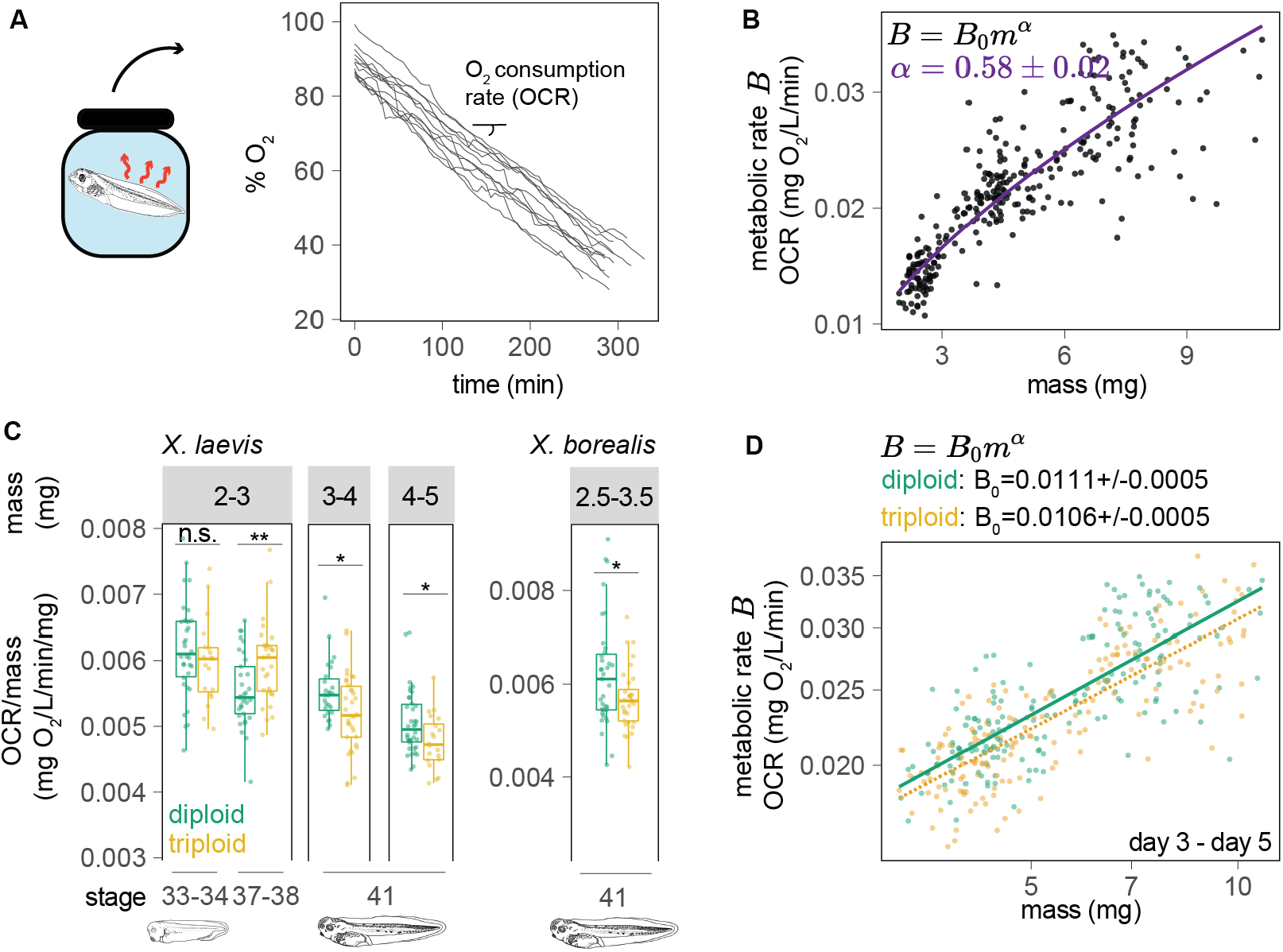
Metabolism assessed by single-tadpole oxygen consumption rates vary between diploids and triploids of both *X. laevis* and *X. borealis*. (**A**) Single tadpoles were placed in individual glass vials sealed from air. The decrease in O2 levels over time was used to calculate the oxygen consumption rate (OCR) for each tadpole. (**B**) OCR as a function of body mass in diploid *X. laevis* embryos from day 2 to 5 post fertilization. The curve shows a fit *B* = *B*_0_*m^α^* with values *B*_0_ = 0.0089 ± 0.0002 and *α* = 0.58 ± 0.02 (n=271) (see SI and Table S2). (**C**) OCR for bins of body mass and stages in *X. laevis* (left panel) and *X. borealis* (right panel) of diploid and triploid embryos (t-test comparing the means, *: p<0.5, **: p<0.01; 3 to 5 clutches per condition, details in Table S3). (**D**) OCR as a function of body mass in diploid (n=189) and triploid (n=165) tadpoles from day 3 (stage 41) to day 5 (stage 46). The lines represent linear fits of the logarithmic values, which yielded the same allometric scaling component exponent α (*α* = 0.46 ± 0.03), but different Y-intercepts that indicate a lower basal metabolic rate (B0) in triploids (see SI and Table S2).

We then compared the metabolic rates of diploid and triploid embryos. To account for the sublinear mass-dependent increase in OCR and potential stage-dependent changes in energy expenditure patterns, measurements were binned by both embryo mass and stage (Fig. 2C, Table S3). At the late tailbud stage 33-34, around 46 hours post fertilization (h.p.f.), we observed no significant difference in mass-normalized OCR. However, a few hours later (~50 h.p.f.) at stage 37-38, triploid embryos displayed a 7.8% higher metabolic rate. This increase was transient and by the early tadpole stage 41 (~72 h.p.f.) triploids displayed a 5.1% lower metabolic rate, a trend that persisted through later developmental stages up to day 5 (Fig. S3A). Due to the ZZ/ZW sex determination mechanism of *X. laevis*, all triploids are female (*24*), and sex-specific differences in metabolic rate have been reported in vertebrates (*25*). However, OCR measurements of male and female diploids at stage 41 showed no significant difference (Fig. S3B). To probe the conservation of this ploidy-dependent OCR decrease, we repeated our measurements using the closely related species *X. borealis* and similarly observed that triploids possessed a lower metabolic rate compared to diploids at stage 41 (8.1%) (Fig. 2C, Fig. S3C). Importantly, plotting OCR vs. body mass for diploid and triploid *X. laevis* tadpoles revealed no difference in the scaling exponent, but rather different normalization constants (Y-intercepts, diploids: *B*_0_ = 0.0111 ± 0.0005, triploids: *B*_0_ = 0.0106 ± 0.0005) (Fig. 2D, Table S2). Therefore, beginning at the tadpole phase, both *X. laevis* and *X. borealis* triploids show a consistent decrease in metabolic rate compared to diploids.

The observed increase in cell size with ploidy at the same body mass indicated that triploid embryos were made of fewer, larger cells than diploids and if so must have decreased rates of cell division, a process previously shown to account for a significant energy cost in zebrafish embryos (*26*). To examine whether differences in cell proliferation explained the variation in metabolic rates, we stained for phospho-histone H3, a marker of mitosis (Fig. 3A), in rapidly growing tails (*27*) (Fig. S4A). Indeed, at the peak of tail growth (stage 41, day 3 p.f.) mitotic cell density was significantly higher in diploid embryos (Fig. 3B), and transient inhibition of cell cycle progression by treatment with palbociclib (*9*), an inhibitor of Cdk4/6, decreased OCR in a dose-dependent fashion (Fig. S4B). However, ploidy-dependent differences in OCR persisted (Fig. 3C) at an inhibitor concentration that significantly reduced proliferation (−85% at the tail base and −72% at the tail tip) and abolished differences in mitotic cell density across ploidies (Fig. S4 C-D). Thus, differences in metabolic rate as a function of ploidy are not explained by altered rates of cell division and the associated energy costs.

**Fig. 3:**
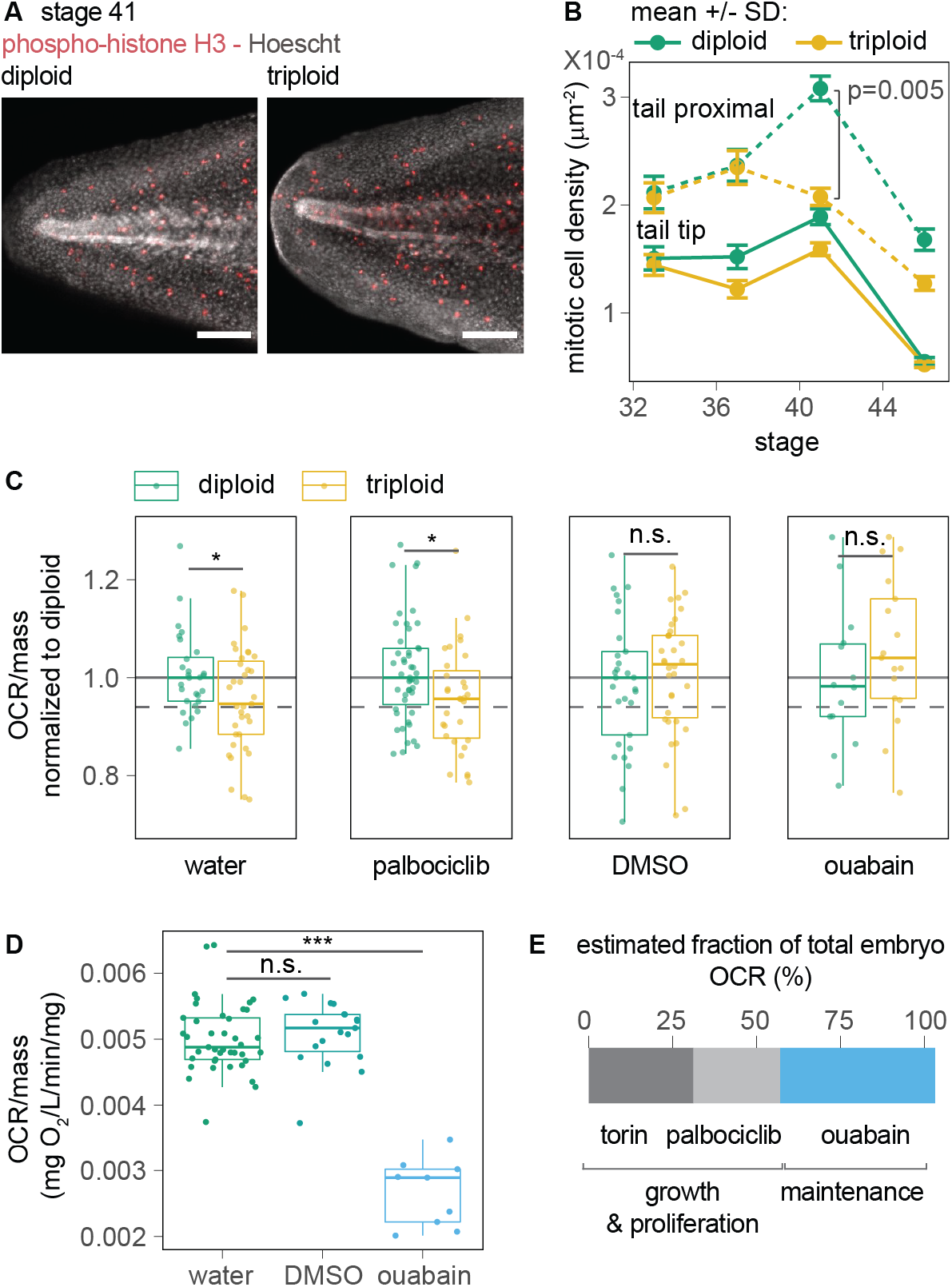
The energetic cost of cell membrane maintenance, not proliferation, accounts for the difference in metabolism between diploids and triploids. (**A**) Representative image of phospho-histone H3 immunostaining and Hoescht DNA dye staining of tadpole tails (scale bar = 200 μm). (**B)**Mitotic cell density assessed by quantifying phospho-histone H3-positive cells (n=3, 5-6 embryos per clutch). (**C**) OCR per mass normalized to the median diploid value for each condition in stage 41 tadpoles during a 6 hr incubation in water, 6 μg/mL palbociclib, 0.1% DMSO, or 200 μg/mL ouabain. Results are shown for 3-4 mg embryos except 5-6 mg for ouabain (n=3 to 5 clutches per condition, details in Table S4). **(D)**OCR per mass for 4-5 mg, stage 41 diploid embryos in water, 0.1% DMSO, or 200 μg/mL ouabain. **(E)**The percentage decrease of OCR in diploids upon treatment with ouabain (Fig. 3D), palbociclib and torin 1 (Fig. S5D) provides an estimate of the contribution of growth and maintenance to the overall embryo energy expenditure. **(B), (C), (D)**, Welch two samples t-test comparing the means, *: p<0.5, **: p<0.01,***: p<0.001.

A second possible explanation for the lower metabolic rate in triploids is a decrease in energy required for cellular maintenance (*21*, *28*, *29*), since processes such as nutrient import and ion gradients depend on cell surface area (*12*, *30*), which scales to the 2/3 power with cell volume. Remarkably, a low dose of DMSO (0.1%) known to perturb membrane stability and ion pumps non-specifically (*31*, *32*) abolished the OCR difference between diploids and triploids, as did treatment with ouabain to inhibit the Na^+^/K^+^ ATPase, a ubiquitous membrane pump essential for maintaining cellular osmolarity and membrane potential (*33*, *34*) (Fig. 3C, Fig. S5A-B and Table S4). Unlike DMSO, ouabain decreased OCR by 46% (Fig. 3D), in agreement with estimates in single-cells (*30*). Comparing the relative effect on OCR of diploids treated with palbociclib, ouabain, and torin-1 allowed us to estimate the energetic costs of proliferation, membrane maintenance and biosynthesis, respectively, and revealed that altogether these processes accounted for most of the embryo energy budget at stage 41 (Fig. E, Fig. S5C-D and S6, SI). Combining the experimentally measured costs with a classical energy partitioning model under the assumption that cellular maintenance scales with surface area, the expected ratio of triploid to diploid metabolic rate of 0.942 ± 0.005 (see SI) was in remarkable agreement with our experimental value of 0.94 ± 0.02 (Fig. S3C). Altogether, these results strongly suggest that the ploidy-dependent difference in metabolic rate observed in *X. laevis* embryos originates from differences in total cell surface area and associated membrane maintenance costs.

Polyploidization increases genome size and drives speciation among amphibians (*15*). We wondered whether, like acute changes in ploidy within a species, evolutionary changes in ploidy across species led to mass-normalized differences in metabolic rates. We recently characterized size metrics in diploid (2N) *X. tropicalis*, allo-tetraploid (4N) *X. laevis* (*35*), and dodecaploid (12N) *X. longipes* (*36*) (Fig. 4A), and showed that genome-to-cell size scaling across species emerged late in development at stage 48 (day 7 p.f.) (*37*) (Fig. 4B and Fig S7A). We measured OCR before and after the onset of cell size scaling to determine whether differences in metabolic rate resulted directly from genome size changes or depended on cell size differences. Whereas *X. tropicalis* were too small in mass to compare until week 5 of development (Fig. S7B), *X. longipes* are similar in size to *X. laevis*. At days 3-4 p.f. (stage 41-45), when cell size did not yet scale with genome size, there was no statistically significant difference in OCR between *X. laevis* and *X. longipes* (Fig. 4C). In contrast, after the onset of cell size scaling, comparison of tadpoles from multiple species at stages with overlapping sizes including 7-13 day old (stage 48-50) *X. longipes*, 6-8 day old (stage 48-50) *X. laevis*, and 26-46 day old (stage 50-52) *X. tropicalis*, revealed a negative trend between metabolic rate and genome size (Fig. 4C, Table S5). This result indicates, at least among closely related *Xenopus* species, the existence of a negative correlation between metabolic rate and genome size that is mediated by cell size scaling (Fig. 4D).

**Fig. 4:**
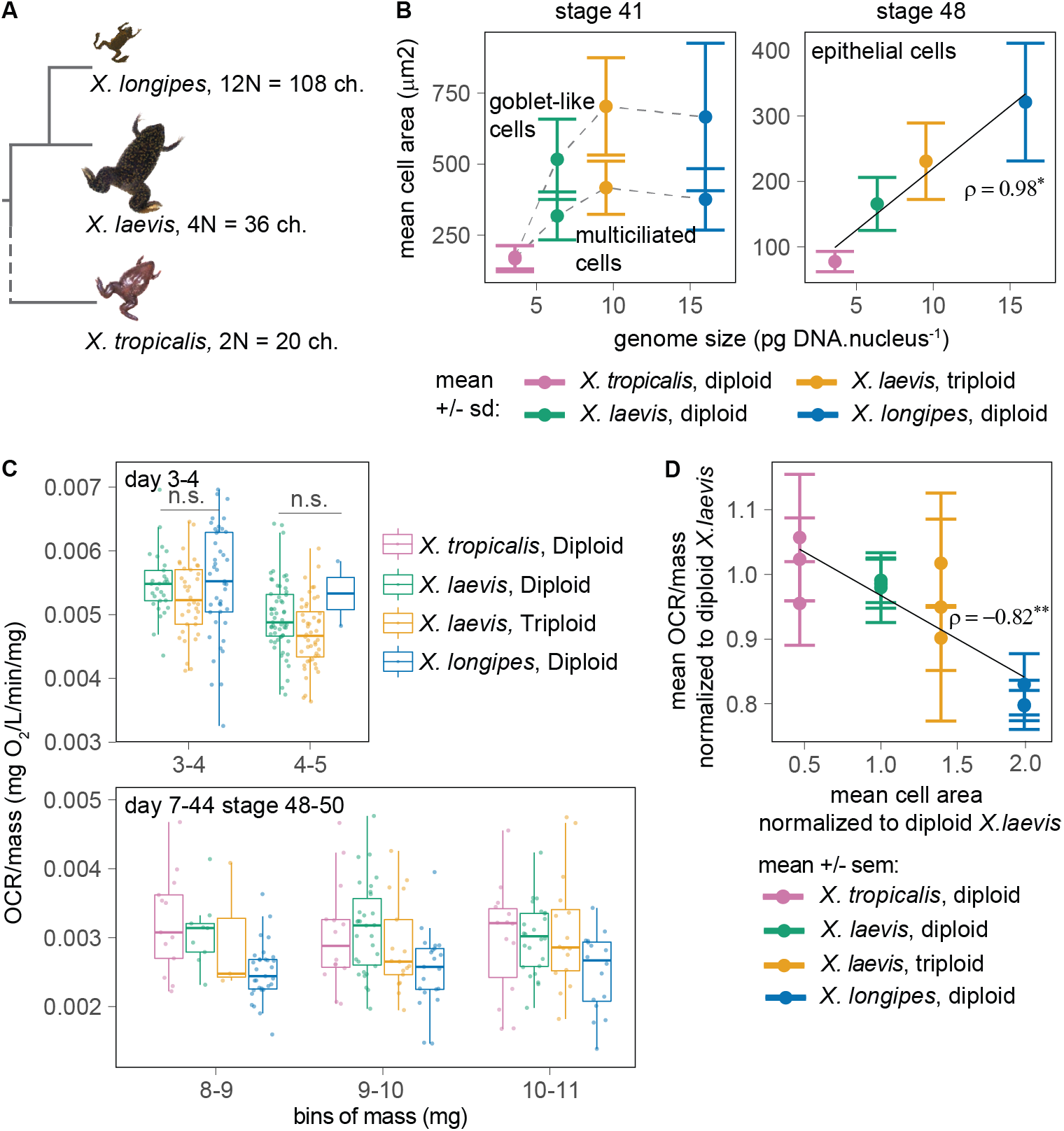
Metabolic rates of *Xenopus* species scale with cell size, not ploidy. (**A**) *X. longipes, X. laevis* and *X. tropicalis* are 12N, 4N and 2N, respectively, with corresponding differences in chromosome (ch.) number; tree not to scale with phylogenetic distances. (**B**) Cell size measurements show scaling with genome size at stage 48, but not stage 41 (2-5 clutches, n>40 per condition, *X. tropicalis* and *X. longipes* stage 48 data from (*37*)). (**C**) OCR across species: (top) before the onset of cell size scaling with genome size at day 3-4 (stage 41-46) and (bottom) after scaling onset (stage 48-50) at day 6-8 (*X. laevis*), 7-13 (*X. longipes*), and 26-46 (*X. tropicalis*) (see Table S5 for n of each bin). **(D)**OCR data at stages 48 or later (from Fig. 4C, bottom) normalized to the mean *X. laevis* diploid value for each bin of mass and plotted as a function of mean cell area at stage 48 (Fig. 4B) normalized to the *X. laevis* diploid value. (B) and (D): ρ is the Pearson’s correlation coefficient of the means weighted by the SD (*: p value <0.05, **: p value <0.01).

In conclusion, this work indicates that increases in cell size driven by polyploidy lower the overall energy expenditure in *Xenopus* larvae by reducing cell maintenance costs. The transient increase in OCR observed in *X. laevis* triploids compared to diploids at stage 37-38 is unexplained, but is concomitant with the emergence of a functional pronephros and may reflect ploidy-dependent adjustments to osmolarity regulation (*38*). We propose that once cell-to-genome size scaling emerges, embryos containing fewer, larger cells and correspondingly lower total cell surface area require fewer active Na^+^/K^+^ pumps and expend less energy maintaining the plasma membrane potential, thereby reducing rates of oxygen and ATP consumption. Thus, the ratio of cell surface area to volume is one factor contributing to whole organism metabolism in vertebrates. While there have been recent advances in theories of metabolic scaling (*39*) and cellular energetics (*40*, *41*), our work establishes a powerful and tractable system to bridge the gap between cell size-dependent energetics and organism metabolism (*12*, *30*, *42*) and address fundamental questions with respect to polyploid tissue metabolism (*1*, *2*, *13*), genome size evolution (*43*), and the embryo energy budget (*26*, *29*).

## Acknowledgments

We thank the Heald lab for feedback throughout the project, the Harland lab, Helen Willsey, Cameron Exner, and Andrea Wills for *Xenopus* embryo protocols, Christian Erikson for help early in the project, Van Savage for helpful conversations, Sarina Arain for her availability with OCR-related questions, Youmna Atieh and Xiao Liu for feedback on the manuscript, James K. Utterback for advice on statistical analysis, and James Evans and the animal facility of UC Berkeley for frog care.

## Funding

National Institutes of Health grant R35GM118183 (RH)

Flora Lamson Hewlett Chair in Biochemistry (RH)

## Author contributions

Conceptualization: CC, RH

Experiments: CC, JB, GO, ML

Analysis: CC

Funding acquisition: RH

Supervision: RH

Writing: CC, RH

## Supplementary Materials

Materials and Methods

Supplementary Text

Figs. S1 to S7

Tables S1 to S5

References (*44-66*)

