## supplementary materials for "Ploidy modulates cell size and metabolic rate in *Xenopus* embryos"

**Cadart et al., 2022**

**This PDF includes:**

- Material and Methods
- Supplementary Text
- Tables S1 to S5
- Fig. S1 to S7
- Additional references

### Materials and Methods

#### Animal care

All frogs were used and maintained following standard protocols established by the UC Berkeley Animal Care and Use Committee and approved in our Animal Use Protocol. Mature *X. laevis, X. tropicalis* and *X. borealis* were obtained from Nasco (Fort Atkinson, WI) or the National *Xenopus* Resource (Woods Hole, MA). *X. longipes* were a kind gift from California Academy of Sciences (San Francisco, CA). All adult frogs were housed in a recirculating tank system with regularly monitored temperature and water quality. *X. laevis* and *X. borealis* were housed at 20-23 ºC, *X. tropicalis* at 23-26 ºC and *X. longipes* at 20-23 ºC.

#### *In vitro* fertilization of *X. laevis, X. tropicalis* and *X. borealis*

The protocol for *in vitro* fertilizations was detailed previously (*44*). For *X. laevis,* females were primed with 100 U of pregnant mare serum gonadotropin (PMSG, National Hormone and Peptide Program, Torrance, CA), at least 48 hr before boosting. The day before the planned experiment (14-16 hr before), females were boosted with 500 U of human Chorionic Gonadotropin (hCG) (Sigma #CG10) and kept at 16 ºC overnight in 1X MMR (1X MMR: 100 mM NaCl, 2 mM KCl, 2 mM $CaCl_2$, 1 mM $MgSO_4$, 0.1 mM EDTA, 5 mM HEPES-NaOH pH 7.6). *X. borealis* females were primed with 60 U of PMSG at least 48 hr before boosting with 300 U hCG 14-16 hr before the fertilization and kept at 16 ºC in dionized water ($diH_2O$). *X. tropicalis* females were primed with 10 U hCG 14-16 hr before priming, kept at room temperature in $diH_2O$ and boosted with 250 U the morning of the experiment. To obtain testes, males were euthanized by over-anesthesia through immersion in 0.05% benzocaine in double-distilled water ($ddH_2O$). Testes were then dissected and kept up to a week at 4 ºC in 1X MR (100 mM NaCl, 1.8 mM KCl, 1 mM $MgCl_2$, 5 mM HEPES-NaOH pH 7.6 in $ddH_2O$) for *X. laevis* and *X. borealis* or placed in a solution of 0.2% BSA (Bovine Serum Albumin) in 1X MBS (1X MBS: 88 mM NaCl, 1.006 mM KCl, 2.49 mM $NaHCO_3$, 0.998 mM $MgSO_4$, 5 mM HEPES-NaOH pH 7.8) and used the same day for *X. tropicalis.* Once they started laying eggs, females were gently squeezed atop a petri dish (60 mm diameter) coated with 1.5% agarose in 1/10X MMR. Each dish of eggs was fertilized with a testis solution obtained by transferring a small piece of testis (~1/4 for *X. laevis* or *X. borealis*, ~1/2 for *X. tropicalis)* in 1 mL $ddH_2O$ *(X. laevis* or *X. borealis)* or 0.5 mL 0.2% BSA in 1X MBS *(X. tropicalis)* in a 1.5 mL Eppendorf tube and crushing it using a plastic pestle. Eggs were gently swirled until they formed a monolayer at the bottom of the dish to ensure that all eggs came into contact with the testis solution, then dishes were slightly tilted to ensure immersion of all eggs and incubated for 10 min (*X. laevis* and *X. borealis)* or 5 min *(X. tropicalis).* After this incubation time, eggs were covered with 1/10X MMR *(X. laevis* and *X. borealis)* or $ddH_2O$ for 10 min, then 1/10X MMR *(X. tropicalis).* To obtain triploid embryos, 1/10X MMR was kept at 4 ºC at least 2 hr before the fertilization, then, exactly 13 min post fertilization (m.p.f.) (*X. laevis)* or 10 m.p.f *(X. borealis),* the media covering the eggs was quickly replaced with ice-cold 1/10X MMR and the dish was placed in a larger dish filled with ice-cold 1/10X MMR in an ice bucket for 15 min (*X. laevis, X.*

*borealis)*. At the end of the cold shock, the media was quickly replaced with room-temperature 1/10X MMR and the petri dish was left to re-equilibrate. To ensure fast temperature-shift of the dishes, petri dishes were not covered with agarose for eggs intended to be triploids. At least 20 m.p.f. (diploid eggs) or 10 min after the cold shock (triploid eggs), fertilized eggs were de-jellied by incubating for ~5-8 min in freshly prepared de-jellying solution (3% L-cysteine in 1/10X MMR-NaOH, pH 7.8). As soon as the jelly coat was dissolved, eggs were washed at least 4 times with 1/10X MMR. At stages 2-3, fertilized eggs were sorted to remove all the unfertilized eggs and incubated at 24 ºC.

#### Natural mating of *X. longipes*

*X. longipes* males and females were injected with a priming dose of 75 U hCG the evening (~4 PM) before the planned mating day and kept at room temperature in separate tanks containing 1 X MMR in di$H_2O$. 10-11 hr later, the morning of the mating day (~3 AM), females were injected with a boosting dose of 200 U hCG. Amplexus typically started 6-8 hr later and eggs were collected in batches approximately every hour. Embryos were de-jellied and kept at 24 ºC as with other *Xenopus* species embryos.

#### Staging of *Xenopus* embryos

Embryos were staged according to the Nieuwkoop and Faber development table (*45*). For stages 41 or older, tadpoles were transiently anesthetized by adding benzocaine to a final concentration of 0.008‰. This allowed manipulation of the embryos by turning them upside down using a hair tool and checking the number of gut loops, which is crucial for staging at stages 41-47. Benzocaine was then rinsed off and tadpoles typically recovered and started swimming again within 10 min.

#### Maintenance of *Xenopus* embryos

On day 1 p.f. (post fertilization), embryos were transferred to larger dishes (100 mm diameter × 15 mm high). Dead or lysed embryos were removed and 1/10X MMR was changed ~2-3 times a day. To avoid potential sources of noise in embryo growth arising from variations in embryo density, *X. laevis* embryos were kept at 60 embryos or tadpoles in 60 mL 1/10X MMR per dish from day 1 onwards. For experiments on late-stage tadpoles in Fig. 4 and S5B-C (day 7 or later), tadpoles were transferred to large containers with 2 L (week 2-3) or 4 L (week 4-7) di$H_2O$ and tanks were kept in the lab at room temperature with a dark/light cycle of ~10/14 hr. Beginning on the morning of day 5, all tadpoles were fed everyday *ad libitum* with the light phase of decanted Sera Micron 5 mg/mL (sera #00720). Sera micron was decanted because the heavier, larger flakes tended to choke young tadpoles.

#### Metaphase spreads

The protocol was adapted from a procedure kindly shared by the Harland lab. To perform metaphase spreads, 10-12 embryos at the tailbud stage (stage 26-34) were euthanized in 0.01‰ benzocaine in 1/10X MMR. The dorsal halves of the tadpoles were then dissected, removing as much of the yolky ventral portion as possible. Dorsal halves were incubated in colchicine (Sigma, #C9754) 1.2 mg/mL in 1/10X MMR for 1.5 hr (the stock was made fresh each time at 40 mg/mL in $ddH_2O$), then incubated in $ddH_2O$ for 20 min, transferred to 0.5 mL Eppendorf tubes and incubated in 200 µL of 60% glacial acetic acid in $ddH_2O$ for 5 min. Each dorsal half was gently transferred with a wide bore 200 µL pipette to a positively-charged glass slide (e.g., Fisherbrand, #12550400). Up to two dorsal halves per glass slides were placed on each glass slide, separated enough that the tissue would not mix once spread out. Excess liquid was blotted away if necessary and a large coverslip (e.g. VWR, 24 × 60 mm, No. 1.5, #48393-251) was put on top of the dorsal halves. To spread the chromosomes, a lead brick was placed on top of the glass slide protected by a paper towel for 5 min. When placing or removing the paper towel and lead brick, extra care was taken to prevent the coverslip from sliding (which otherwise would give metaphase spreads that were hard to quantify). The glass slide and coverslip were then put on dry ice for 5 min. Quickly after removal from the dry ice, the coverslip was detached from the glass slide using a razor blade and gently prying up the edge of the coverslip from the frozen slide. Once the coverslip was removed, the metaphase spread was left to dry for 1-3 min. Once dry, the slide was mounted by putting a drop of Vectashield antifade mounting medium (Vector Laboratories, #H-1000-10) with Hoescht 0.1 mg/mL, covering with a coverslip (e.g. Thomas Brand, 22 × 22 mm No. 1.5 #1169W10) and sealing with nail polish. Slides were then stored in a box protected from light at 4 °C until imaging.

#### Mass measurements

Mass measurements were performed using the Mettler Toledo Excellence XSR Analytical Balance with a 0.01 mg readability (#XSR105DU). One by one, each tadpole was transferred onto a glass coverslip, excess water was removed using a pipette and blotting with kimwipes (Kimberley-Clark #34120). Rapidly after the measurement, the tadpole was placed back into water and, if needed, used for further analyses (e.g., proliferation assay after palbociclib treatment, Fig. S4C-D).

#### Survival and developmental curves

For the survival and developmental curves (Fig. S1A-B), three 60-mm diameter dishes with 30 embryos per ploidy and clutch were made on day 0 and survival was recorded 2 times per day during the first 3 days, then once per day for another 4 days (Fig. S1A). For the developmental curve, to minimize the effect of temperature fluctuations on developmental rate, all dishes were placed on the middle shelf of the incubator and a rotation was established among the three dishes for each condition (*46*): at each time point, the dish at the front of the shelf was taken out to assess the average stage in the dish, then placed at the back of the incubator, and so forth.

**Oxygen consumption rate (OCR) measurements**

Oxygen consumption rate measurements were performed using two Presens Sensor Dish Reader (SDR) v4 sets (Presens, #200001059) and 1 mL (day 1-2 p.f. embryos), 2 mL (Presens, #200001798) (day 3-5 p.f.) or 4 mL (Presens, #200001369) (day 7 or later p.f.) SensorVials which allowed measurement of up to 48 samples per experiment. For all the experiments, the system was placed in an incubator at 24 ºC with no light and measurements were made with one embryo per vial in dd$H_2O$. The day before the experiment, the SensorVials and two to four 100 mL bottles containing ~60 mL of dd$H_2O$ were placed in a second 24 ºC incubator for temperature equilibration. The morning of the experiment, 24 diploid and 24 triploid embryos were staged and transferred into a large dish containing dd$H_2O$ to rinse off the 1/10X MMR solution. To oxygenate the water, the bottles with dd$H_2O$ were shaken vigorously for 60 sec, then gently tapped onto a table to help bring the bubbles to the surface. It was crucial not to introduce any bubbles in the vial. One by one, embryos were then transferred into a vial, the vial was filled to the brim with water, making sure that no bubbles were introduced, and the vial was sealed tightly with the cap and placed onto a transparent-bottom 24 multi-well plate. The incubator containing the SDR reader was opened only once, just briefly, to place the two multi-well plates and it was kept tightly closed for the rest of the experiment. $O_2$ levels were recorded every 5 min for 5-7 hr using the Presens software.

For each experiment, one of the vials contained no embryos in order to assess the time at which $O_2$ levels became stable (once the temperature had equilibrated). All timepoints acquired prior to this time were removed from the analysis (typically the first 2-3 hours, Fig. S2A). Within each experiment, $O_2$ concentration in the blank vial fluctuated less than 0.4 ± 0.2% on average (Fig. S2B) and temperature fluctuated less than 0.13 ± 0.06% on average (Fig. S2C), thus indicating that measurements were made at near-equilibrium with negligible temperature fluctuations. All experiments were performed in incubators set at 24 ºC and the temperature measured by the SDR device was on average 23.4ºC, with only 0.92% fluctuations across all 66 experiments used for Fig. 2B-D and 4 C-D (Fig. S2D).

To calculate the rate of consumption of $O_2$ for each tadpole, a custom program in R was written to allow the user to select, for each curve, the first and last point between which to calculate a robust linear fit, using the 'lmrob' function with the MM-estimator algorithm (*47*) from the 'robustbase' package in R. During this process, the user could also exclude from the analysis any curve which presented too much noise (e.g., in case an air bubble caused a sudden drop in the curve). All the fits obtained were performed over at least 2 hr of continuous $O_2$ recording (Fig. S2E), yielded R2 values that were higher than 0.90 (Fig. S2F) and the calculated slopes (rates) were all statistically significant with p values smaller than 0.001 (Fig. S2G).

Given the high variability in the OCR measured across individual tadpoles, we established the following controls and procedures to ensure robust results:

- Once we begun identifying a decrease in OCR in triploids compared with diploids at stage 41 (Fig. 2C), we further confirmed the result by performing 3 blinded experiments in which the diploid and triploid dishes were randomly assigned a letter by another lab member. The ploidy was kept secret from the experimentalist until after the entire experiment was over, including tadpole staging, OCR recording, and data plotting and statistical analysis.

- During the analysis, the step at which the user could choose to remove a curve if it showed too much noise or abnormal variations or if the quality of the linear fit seemed unconvincing was done prior to matching the curve with the sample name and ploidy.
- A second step in the R analysis program, also prior to matching the measurement with the ploidy of the embryo, presented a plot showing OCR *vs*. embryo mass. This allowed identifying clear outliers (i.e., because of a mistake in the wet mass measurement or because of a problem on the linear fit which was always checked a second time in this case). However, most often, no outliers were identified at this step.
- All our results were obtained by comparing at least 3 clutches containing diploid and triploid embryos. The details of the number of clutches and individual embryos measured for each OCR result are detailed in Tables S3-5.

Occasionally, a tadpole died during the measurement, halting a decrease of $O_2$ in the vial (Fig. S2H). The absence of decrease was a clear indication that the potential contribution of bacteria to the measured oxygen consumption was negligible compared with that of embryos. Tadpoles that died during the experiment were excluded from the analysis.

**PCR for sex determination**

The protocol for PCR-based sex determination was kindly shared by Helen Willsey's lab (UCSF, CA) and was adapted from ref. (*24*). Following OCR measurement, tadpoles were euthanized with 0.01‰ benzocaine in dd$H_2O$ and the tail of each tadpole was sectioned, transferred to a 200 µL PCR tube, spun down to remove excess water and flash frozen at –80 ºC. To lyse the tails, 20 µL of 1X Phusion buffer (New England Biolabs, #E0553S) was added to each tube, tails were spun down briefly to ensure that they were submerged and the following lysis program was started: boil at 95 ºC for 10 min, add 2.5 µL of 20 ng/mL Proteinase K (New England Biolabs, #P8107S) to each well, incubate at 55 ºC for 4 hr, boil again at 95 ºC for 10 min, store at 4 ºC. The PCR reaction was performed using the Phusion High-Fidelity PCR kit (New England Biolabs #E0553S) following the manufacturer's instructions. The following parameters were used: denaturation: 95 ºC, 30 sec; initiation: 98 ºC 10 sec; melting: 58 ºC, 30 sec; elongation: 72 ºC, 30 sec; extension: 72 ºC 10 min. Forward (AAAACCATGACCTCCCGGATAC) and reverse (TAGGGAGGGGTTTGGAGGTTC) primers amplified a 315 base-pair amplicon from the gene W3 whose presence is indicative of a female genotype. 100% of the triploids genotyped were positive for the gene W3 while the ratio of W3 positive tails in diploids were 11/23, 10/23 and 13/23 for the three clutches analyzed.

**Phalloidin staining and immunofluorescence**

The protocol for immunofluorescence combined with phalloidin staining was obtained from Helen Willsey's lab (UCSF, CA). The protocol for phospho-histone H3 staining was inspired by ref. (*27*).

6-10 embryos of the desired stage were collected for each ploidy, fixed for 45 min in 4% PFA in PBS, then rinsed several times in PBS and kept at 4 ºC until use. Embryos were then permeabilized by incubating 3 × 20 min in PBS with 0.01% Triton x-100 (PBT), blocked for 1 hr at room temperature with PBT containing 10% goat serum. For phospho-histone H3 (pH3) staining for the tail proliferation assays, the following steps were performed: embryos were (a) incubated overnight at 4 ºC with 1:1000 mouse anti-histone H3 (phosphor Ser10), Abcam 14955, (b) rinsed 3 × 20 min in PBT, (c) incubated overnight at 4 ºC with 1:500 of the secondary antibody (goat-anti mouse coupled with AlexaFluor 488, Invitrogen), (d) rinsed again 3 × 10 min in PBS, (e) incubated with 0.01 mg/mL Hoechst for 20 min, (f) rinsed briefly with PBS, (g) mounted on slides in Vectashield just before imaging.

For cell size measurements, TMR-phalloidin 1X (Thermo Scientific #R415, stock at 400 X in DMSO according to the manufacturer's instructions) was added at step (3) or the experiment was done without steps (1)-(2). For cell size measurements at stage 47 (Fig. 4 B), phalloidin staining could not be used because it overwhelmingly stains the thick F-actin bundles along the tail muscle. We thus turned to immunofluorescence staining of ZO-1 to stain the cell membrane (Thermo Scientific #33-9100). The following protocol, adapted from ref. (*48*), was used. First, tadpoles were fixed for 1 hour in MAD fixative (2 parts methanol, 2 parts acetone, 1 part DMSO) then dehydrated in methanol and stored at 20 ºC. On day 1 of the staining procedure, the samples were: (a) gradually rehydrated in 0.5X SSC (1X SSC: 150 mM NaCl, 15 mM sodium citrate, pH 7), (b) bleached in bleaching solution (0.5X SSC solution containing 2%, $H_2O_2$ and 5% formamide) for 2-3 hr under light, (c) washed 3 × 10 min in PBT-BSA (PBT containing 2 mg/mL BSA), (d) blocked in PBT-BSA supplemented with 10 % goat serum and 5 % DMSO for 1-3 hr and finally (e) incubated at 4 ºC overnight in 300 µL of primary antibodies (ZO-1 at 1:250). The second day of the staining, samples were rinsed 4 × 2 hr in PBT then incubated overnight in PBT supplemented with 1:500 goat anti-mouse secondary antibody coupled with Alexa Fluor 488. On day 3, tadpoles were washed 4 × 2 hr in PBT and gradually dehydrated in methanol. Embryos were then cleared in Murray's clearing medium (2 parts Benzyl Benzoate, 1 part Benzyl Alcohol).

#### Drug treatments

For each drug, the working concentration was determined in two steps. First, 5 embryos were incubated in 10 mL of increasing doses of each drug for 6 hours to assess the concentration which killed the tadpoles. Working with a narrower range of concentrations below the lethal one, we then performed dose–response measurements of OCR (Fig. S4E, S5A and S6A) and selected the lowest concentration at which OCR decrease was maximal and beyond which it decreased very little.

Palbociclib stock (LC Laboratories #P-7744) was made at 50 mg/mL in $ddH_2O$-HCl pH 1 and stored at –20 ºC (*49*). It was then diluted in $ddH_2O$ to a final concentration of 6 µg/mL (Fig. 3C and S4C-D), unless specified (Fig. S4B). Ouabain stock was made fresh at 5 mg/mL in $ddH_2O$ and diluted in $ddH_2O$ to a final concentration of 200 µg/mL (Fig. 3C-D), unless specified for the dose-response curve (Fig. S5A). Torin-1 stock was made at 10 mg/mL in DMSO and stored at –20 ºC, then diluted to a final concentration of 1 μg/mL (final DMSO concentration was 0.1%) unless specified for the dose-response curve (Fig. S6A).

**Tail length measurements**

Stereoscope images were obtained on a Zeiss® Stereoscope Discovery.V8 with the Leica color camera DNC 2900 and LASX software. Embryos stage 41 or older were transiently immobilized with a low dose of benzocaine (0.008‰ in 1/10X MMR) and placed on 1.5% agarose-coated petri dishes for imaging. Images were analyzed using Fiji software.

**Microscopy imaging**

Confocal imaging was performed on an inverted Zeiss LSM 800 using the Zeiss Zen software, a Plan-Apochromat 20X/0.8 air objective with a pinhole size set at 1 Airy Unit (cell size measurement) or Plan-Apochromat 10X/0.45 air objective with a pinhole opened to the maximum diameter (phospho-histone H 3 imaging for tail proliferation assays). For the metaphase spreads, images were acquired using Micromanager 1.4 software (*50*) on an upright Olympus® BX51 microscope equipped with an ORCA-ER and ORCA-Spark camera (Hamamatsu, Photonics) and Olympus UPlan 60X/NA 1.42 oil objective.

**Cell size measurement**

Cell areas were manually traced using Fiji software. For each condition, at least 3 clutches and 3-5 embryos per clutch and ploidy were measured.

**Statistical analysis**

All the analysis and figures were made using R 4.1.2 and the following packages: ggplot2 (*51*), tidyverse (*52*), dplyr (*53*), forcats, robustbase and MASS. T-tests in Fig. 1D, 2B, 2D, 2F are Welch two samples t-tests.

### Supplementary text

#### Testing the effect of sex on OCR

Since all triploids are females, we checked that the difference in OCR as a function of ploidy did not arise from an imbalance in the sex ratios of embryos. We performed OCR measurements on 3 clutches of 23 stage 41 diploid embryos, collected their tails and performed PCR for sex determination, then matched the sex genotype of each tadpole to the OCR measured. We then plotted OCR in diploids as a function of the sex of the embryo (Fig. S3B). We found no statistically significant difference, indicating that this was not a cause of OCR decrease in triploids.

#### Testing the existence of a power-law relationship between OCR and body mass in embryos

To test the existence of a power-law relationship between OCR and body mass, as reported in adult animals according to Kleiber's law (*20*), we first focused on diploid *X. laevis* embryos during day 2-5 p.f. which offered a significant range of body mass (Fig. 2B). We performed linear regressions on the logarithmic or non-transformed values of OCR and body mass. The former gave a higher $R^2$ (Table S2), indicating that the data was better represented by a power-law relationship of the form: $B = B_0 m^{\alpha}$ (where $B$ is OCR and $m$ is body mass) than by a simple linear relationship (where $B = am + b$). In Fig. 2D, we compared this scaling relationship in diploids and triploids from day 3-onwards, because the relationship between OCR and ploidy changes during day 2 (Fig. 2C). While the range of body masses is narrower and the allometric exponent changes a little compared with the one obtained with the full diploid dataset (day 2-5) this analysis shows distinct normalization constant $B_0$ as a function of ploidy (Table S2).

#### Note on the interpretation of the effects of torin-1 on OCR

To test the effect of inhibiting biosynthesis, we measured OCR in tadpoles treated with torin-1, a specific inhibitor of the mTOR pathway (*54*, *55*). While *X. laevis* are tolerant to a range of DMSO concentrations, up to 0.1% (*56*, *57*), we found that this concentration abolished the difference in OCR between diploid and triploid embryos (Fig. 3C) due to its interactions with the cellular membrane (*31*, *32*). Unfortunately, torin-1 could only be reconstituted in DMSO, which compromised the assessment of its effect on diploids compared to triploids (Fig. S6C). Upon treatment with torin-1, the difference in OCR between diploid and triploids was qualitatively conserved for 3-4 mg embryos (Fig. S6B, left) and lost for 4-5 mg embryos (Fig. S6B, right), but it is impossible to parse out the contributions of DMSO from that of torin-1 in this result. However, since DMSO treatment does not impact the average OCR value of diploids (Fig. 3D), it is possible to estimate the decrease of OCR upon biosynthesis inhibition with torin-1 (Fig. S6A), which is about ~30% of the OCR of a diploid embryo (Fig. 3E).

**Model for the relationship between metabolism and cellular maintenance**

Previous theoretical work proposed partitioning organism metabolism ($B$) into two main processes, growth and maintenance (*13*, *28*, *58*), a model known as the Von Bertalanffy-Richards model of growth (*59*, *60*). Here we propose a slight adaptation of this model to estimate costs of growth and maintenance as a function of ploidy in embryos.

First, we write that:

$$B = \frac{E_S}{m_c}\frac{dm}{dt} + E_d\frac{dN_c}{dt} + B_c N_c \qquad (1)$$

where $E_s$ is the cost of synthesis of new units of mass $m$, $m_c$ is the mass of a cell, $E_d$ is the cost of new cell division, $\frac{dN_c}{dt}$ is the rate of cell proliferation, $B_c$ is the cellular metabolic rate, and $N_c$ the number of cells in the embryos. Note that, compared with previous work in multicellular tissues (*13*, *28*, *58*–*60*), we propose to partition the term "growth" into two distinct processes: biosynthesis and cell cycle progression. Our experimental results however suggest that neither process scales differently with ploidy (Fig. 3C), so we are left to estimate how differences in maintenance affect the overall metabolic rate.

How does cellular maintenance change with ploidy? A first simple hypothesis is that cellular metabolic rate scales linearly with cell volume. Since cell size measurements suggest a linear scaling between ploidy ($P$) and cell volume ($V_c$) in epithelial cells (Fig. 1E-F), we can estimate that for an embryo volume $V_{\text{embryo}}$, the number of cells follows:

$$N_c \sim V_{embryo}/V_c \sim 1/P \qquad (2)$$

In this case, the product $B_c N_c$ remains constant with ploidy, a result which disagrees with our experimental measurements that OCR decreases with ploidy (Fig. 2C). In the alternative hypothesis where cellular metabolic rate remains constant with cell volume, we instead find: $B_c N_c \sim 1/P$.

Finally, a third hypothesis is that cellular maintenance scales with cell surface area, rather than volume. This is supported by our finding that perturbing membrane stability and ion pumps using DMSO and ouabain abolishes the OCR difference between diploid and triploids (Fig. 3C). Importantly, previous work in yeast (*6*), human stem cells (*61*), and unicellular organisms (*62*) also suggested that cellular respiration or metabolic rate increases sub-linearly with cell size (reviewed in (*12*)). Assuming a scaling with surface area, we find:

$$B_c N_c \sim P^{-1/3} \qquad (3)$$

To estimate the fraction of the total energy expenditure associated with cell maintenance ($\varphi_m$), we turn to our experimental measurements of OCR upon inhibition of biosynthesis (torin-1), cell cycle progression (palbociclib) and membrane potential maintenance (ouabain) which suggest that these processes represent 31 ± 5 %, 26 ± 1 %, and 46 ± 4 % of total embryo energy expenditure respectively (Fig. 3E). Note that the sum of these contribution is 103 ± 6 % (variation calculated by standard error propagation). With the margin of error, the sum is thus in agreement with 100%, which suggests that these three processes account for the major part of all relevant energetic costs. Moreover, the estimated cost of membrane potential maintenance by the $Na^+/K^+$ ATPase is within the range of previously published values (*30*, *63*, *64*).

To estimate the metabolic rate of diploids ($B_D$), we use equation (1) and write:

$$B_D = (1 - \varphi_m)B_D + B_c N_c \qquad (4)$$

Under the experimentally tested assumption that growth and maintenance are constant with ploidy, the metabolic rate of a triploid embryo is thus:

$$B_T = (1 - \varphi_m)B_D + (B_c N_c)_T \qquad (4)$$

Under the assumption that cellular metabolic rate scales with surface area as in equation (3),

$$(B_c N_c)_T = P^{-1/3}(B_c N_c)_D = P^{-1/3}\varphi_m B_D \qquad (5)$$

Combining equations (4) and (5) and using the experimentally measured $\varphi_m$ = 46 ± 4 %, the estimated ratio of metabolic rate of triploids over diploids is $\mathrm{B_T}/\mathrm{B_D} = 0.942 \pm 0.005$. This is in remarkable agreement with our average experimentally measured ratio of $0.94 \pm 0.02$ across *X. laevis* and *X. borealis* (Fig. S3C).

We note that it is possible that some experimentally measured costs might be overestimated due to overlaps between the effect of the drugs; inhibition of biosynthesis, for example, is likely to alter cell cycle progression because of feedback between the two pathways (*65*) while inhibition of the $Na^+/K^+$ ATPase by ouabain alters osmolarity in addition to plasma membrane potential (*33*, *34*), which can have multiple impacts on cellular physiology including biosynthesis (*11*, *34*, *66*). These considerations could contribute to the uncertainty of the above estimate.

| stage | N | nD | nT | pval |
|---|---|---|---|---|
| 33-34 | 3 | 33 | 26 | 0.54 |
| 35-36 | 1 | 10 | 10 | 0.71 |
| 37-38 | 5 | 49 | 50 | 0.19 |
| 41 | 5 | 82 | 64 | 0.0031 |
| 43 | 3 | 33 | 31 | 0.025 |
| 45-46 | 5 | 66 | 61 | 0.47 |
| 47 | 3 | 34 | 33 | 0.29 |

**Table S1. Details of experiments used and statistics for mass measurements in Fig. 1D.** N is the number of different clutches, nD the number of diploids, nT the number of triploids and pval indicates the p value from a pairwise t-test comparing the means across ploidies for a given stage.

| days | P | α | SE of α | $B_0$ | SE $B_0$ | $R^2$ power law model | $R^2$ linear model | N | n |
|---|---|---|---|---|---|---|---|---|---|
| day 2-5 | D | 0.58 | 0.02 | 0.0089 | 0.0002 | 0.83 | 0.76 | 19 | 271 |
| day 3-5 | D | 0.46 | 0.03 | 0.0111 | 0.0005 | 0.59 | 0.55 | 11 | 189 |
| day 3-5 | T | 0.46 | 0.03 | 0.0106 | 0.0005 | 0.66 | 0.645 | 11 | 165 |

**Table S2. Results of power-law fits for Fig. 2B and 2D.** Measurements made on *X. laevis* embryos. P is the ploidy. The table reports the calculated allometric scaling exponent α and its standard error (SE), the calculated normalization constant of the power-law $B_0$ and its standard error. The $R^2$ values for a linear fit on the logarithm-transformed ($R^2$ log) and non-transformed ($R^2$ linear) data are shown for comparison. N is the number of independent clutches, n is the number of individual embryos measured.

| species | stage | bin of mass (mg) | N | nD | nT | mean OCR/mass diploid (mg $O_2$/L/min/mg) | mean OCR/mass triploid (mg $O_2$/L/min/mg) | pval |
|---|---|---|---|---|---|---|---|---|
| *X. borealis* | 41 | 2.5-3.5 | 3 | 35 | 32 | 0.0062 | 0.0057 | 0.021 |
| *X. laevis* | 33-34 | 2-3 | 3 | 34 | 19 | 0.0062 | 0.0060 | 0.34 |
| *X. laevis* | 37-38 | 2-3 | 4 | 33 | 29 | 0.0055 | 0.0060 | 0.0067 |
| *X. laevis* | 41 | 3-4 | 5 | 27 | 34 | 0.0056 | 0.0052 | 0.013 |
| *X. laevis* | 41 | 4-5 | 5 | 38 | 20 | 0.0051 | 0.0048 | 0.019 |
| *X. laevis* | 43 | 4-5 | 3 | 17 | 25 | 0.0049 | 0.0046 | 0.18 |
| *X. laevis* | 45-46 | 6-7 | 3 | 11 | 13 | 0.0042 | 0.0042 | 0.62 |

**Table S3. Details of experiments used and statistics for OCR measurements in Fig. 2C and Fig. S3A.** N is the number of independent clutches, nD the number of diploids, nT the number of triploids, and pval indicates the p value from a pairwise t.test comparing the means across ploidies for a given stage and size of bin. Values are calculated only for conditions with more than 10 observations but Fig. S3A shows all the bins.

| treatment | bin of mass (mg) | median D | median T | SD D | SD T | pval | ratio | nD | nT | N |
|---|---|---|---|---|---|---|---|---|---|---|
| water | 3-4 | 1 | 0.95 | 0.085 | 0.11 | 0.0384 | 0.95 | 28 | 35 | 5 |
| water | 4-5 | 1 | 0.97 | 0.11 | 0.085 | 0.0404 | 0.97 | 41 | 21 | 5 |
| DMSO | 3-4 | 1 | 1 | 0.15 | 0.12 | 0.869 | 1 | 30 | 34 | 6 |
| DMSO | 4-5 | 1 | 0.98 | 0.074 | 0.15 | 0.584 | 0.98 | 19 | 26 | 5 |
| torin-1 | 3-4 | 1 | 0.95 | 0.13 | 0.12 | 0.797 | 0.95 | 15 | 8 | 2 |
| torin-1 | 4-5 | 1 | 1 | 0.15 | 0.17 | 0.391 | 1 | 18 | 25 | 4 |
| torin-1 | 5-6 | 1 | 1.2 | | 0.24 | | 1.2 | 1 | 2 | 1 |
| palbociclib | 3-4 | 1 | 0.96 | 0.1 | 0.11 | 0.0203 | 0.96 | 47 | 31 | 3 |
| palbociclib | 4-5 | 1 | 1.2 | 0.12 | 0.085 | 0.0063 | 1.2 | 10 | 9 | 1 |
| ouabain | 4-5 | 1 | 1.4 | 0.22 | 0.19 | 0.0813 | 1.4 | 4 | 5 | 2 |
| ouabain | 5-6 | 1 | 1.1 | 0.23 | 0.51 | 0.268 | 1.1 | 17 | 20 | 4 |

**Table S4. Details of experiments used and statistics for OCR measurements in Fig. 3B.** OCR values were normalized to the median diploid value for each bin and drug treatment. Median D and T are the median OCR per mass values for diploid and triploids, respectively, after normalization, SD D and T are the standard deviations of OCR per mass for diploids and triploids, respectively. Ratio is the ratio of median OCR of triploids over diploids. N is the number of independent clutches, nD the number of diploids, nT the number of triploids, and pval indicates the p value from a pairwise t-test comparing the means across ploidies for a given stage and size of bin.

| bins of mass (mg) | species | ploidy | mean OCR/mass (mg $O_2$/L/min/mg) | SD OCR/mass (mg $O_2$/L/min/mg) | n | N | day |
|---|---|---|---|---|---|---|---|
| 3-4 | *X. laevis* | D | 0.10 | 0.09 | 29 | 6 | 3 |
| 3-4 | *X. laevis* | T | 0.96 | 0.11 | 36 | 6 | 3 |
| 3-4 | *X. longipes* | D | 1.00 | 0.18 | 45 | 3 | 4 |
| 4-5 | *X. laevis* | D | 1.02 | 0.11 | 59 | 8 | 3 |
| 4-5 | *X. laevis* | T | 0.96 | 0.11 | 48 | 8 | 3-4 |
| 4-5 | *X. longipes* | D | 1.09 | 0.15 | 2 | 2 | 4 |
| 8-9 | *X. laevis* | D | 0.98 | 0.16 | 9 | 5 | 7 |
| 8-9 | *X. laevis* | T | 0.95 | 0.31 | 3 | 1 | 7 |
| 8-9 | *X. longipes* | D | 0.80 | 0.13 | 31 | 3 | 7-13 |
| 8-9 | *X. tropicalis* | D | 1.02 | 0.23 | 13 | 2 | 26-42 |
| 9-10 | *X. laevis* | D | 0.99 | 0.21 | 30 | 5 | 7 |
| 9-10 | *X. laevis* | T | 0.90 | 0.21 | 17 | 4 | 7 |
| 9-10 | *X. longipes* | D | 0.80 | 0.17 | 21 | 3 | 7-13 |
| 9-10 | *X. tropicalis* | D | 0.96 | 0.24 | 14 | 3 | 23-42 |
| 10-11 | *X. laevis* | D | 0.99 | 0.10 | 24 | 6 | 6-7 |
| 10-11 | *X. laevis* | T | 1.02 | 0.27 | 16 | 5 | 6-7 |
| 10-11 | *X. longipes* | D | 0.83 | 0.19 | 16 | 3 | 9-13 |
| 10-11 | *X. tropicalis* | D | 1.06 | 0.38 | 15 | 4 | 22-44 |
| 11-12 | *X. laevis* | D | 1.90 | 0.25 | 9 | 4 | 6-8 |
| 11-12 | *X. laevis* | T | 1.17 | 0.20 | 15 | 4 | 6-8 |
| 11-12 | *X. longipes* | D | 1.004 | 0.21 | 10 | 3 | 9-13 |
| 11-12 | *X. tropicalis* | D | 1.54 | 0.54 | 7 | 3 | 32-46 |

**Table S5. Details of experiments used and statistics for OCR measurements in Fig. 4C.** N is the number of independent clutches, n is the number of individual tadpoles.

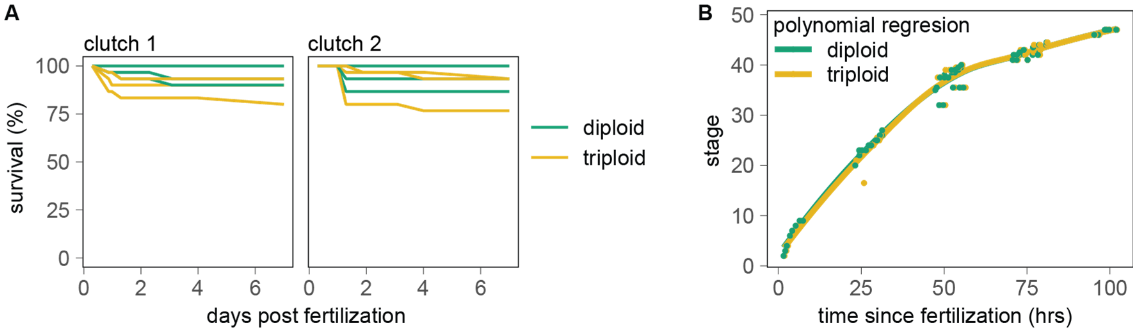


**Fig. S1. Triploid embryos develop and survive at rates comparable to diploids. (A)** Survival rate over the first 7 days post fertilization for two representative clutches. Each line represents a technical replicate. **(B)** Developmental curves of diploids and triploids at 24 ºC (three clutches containing diploids and triploids). The line represents a polynomial fit.

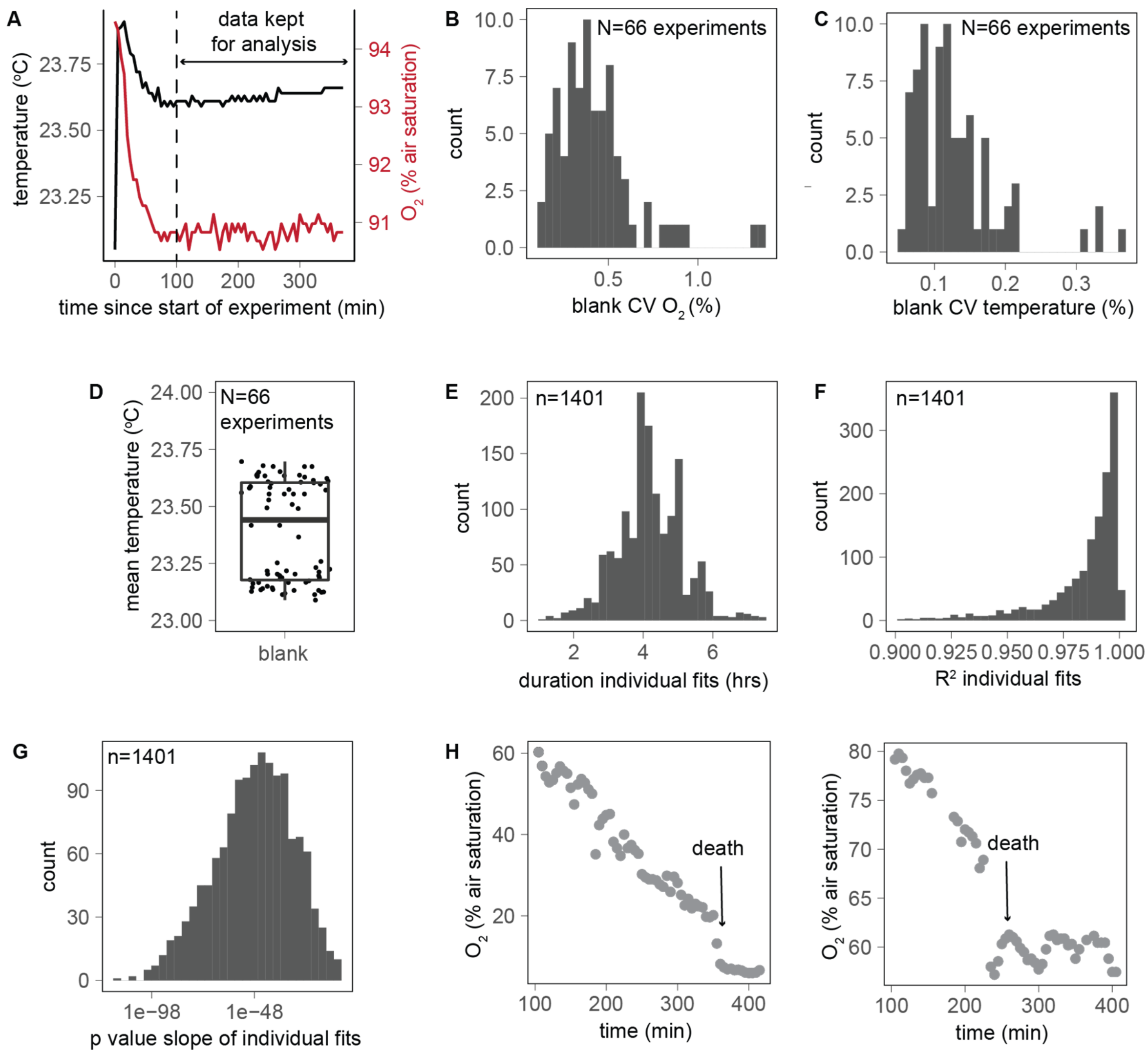


**Fig. S2. Quality controls for OCR experiments from Fig. 2 and 4. (A)** Example of measurements obtained in a blank vial containing no tadpoles. Blanks were used to assess, for each experiment, the time at which $O_2$ and temperature stabilized. Timepoints acquired before equilibrium were discarded. Details of the coefficient of variation (CV) of $O_2$ **(B)** and temperature (**C)** in the blanks of each of the 66 experiments used for figures 2B-D and 4C-D show that these variables were very stable once the timepoints prior to equilibrium were removed. **(D)** Average temperature showed little variation across all experiments. Details of the duration **(E),** $R^2$ **(F)**, and p values (**G)** of all 1401 individual linear fits performed to calculate single-embryo or single-tadpole OCR for all 66 experiments used for figures 2B-D and 4C-D. **(H)** Examples of two OCR experiments during which the embryo died show that there is no $O_2$ consumption after the embryo death, suggesting negligible contribution from potential bacteria to the OCR measurements.

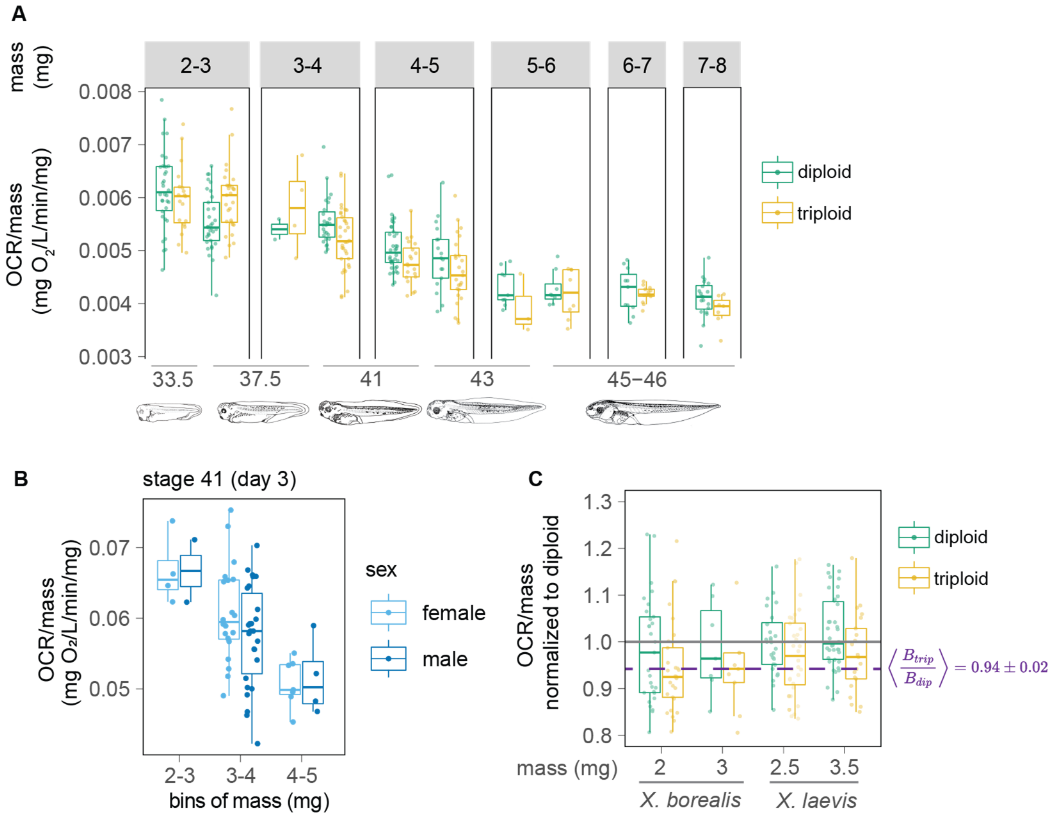


**Fig. S3. Details of OCR measurements.** (**A**) Comparison of OCR normalized to mass and binned by mass and stage over 5 days of development reveals a transient increase followed by consistent decrease in metabolic rate in *X. laevis* triploids compared to diploids (3 to 5 clutches per condition, details in Table S3). (**B**) Comparison of male and female *X. laevis* diploids reveals no sex-related metabolic differences. (**C**) Normalization to diploid animals indicates that both *X. borealis* and *X. laevis* have a ~6% decrease in metabolic rate.

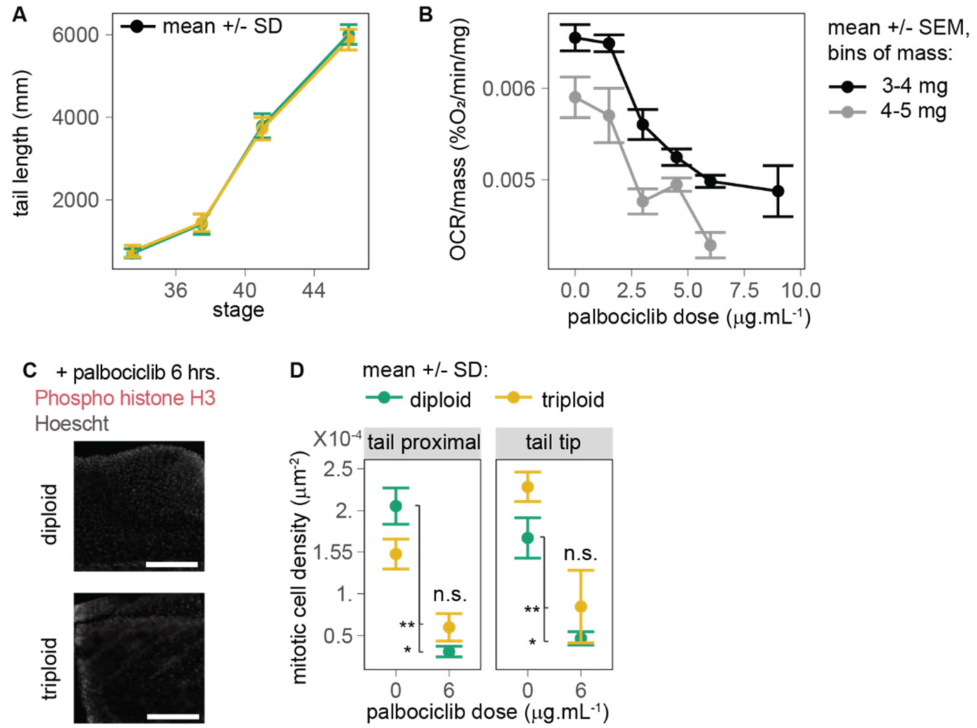


**Fig. S4. Details of proliferation quantification and inhibition experiments. (A)** Tail length throughout development (2-4 clutches per stage, n > 30 embryos per ploidy and stage). **(B)** OCR per mass as a function of palbociclib doses for stage 41 diploid embryos and two bins of mass (6 clutches, n = 2 to 57 per bin of mass and palbociclib concentration). **(C)** Representative image of phospho-histone H3 and Hoescht staining in tails as in Fig. 3A in diploid and triploids following a 6-hr treatment with 6 µg/mL palbociclib (scale bar = 200 µm). **(D)** Mitotic cell density in tails after 6 hr treatment with or without 6 µg/mL palbociclib (2 clutches, 5-6 embryos per clutch and condition; Welch t-test comparing the mean, *: p<0.05, **: p<0.01, n.s.: not significant).

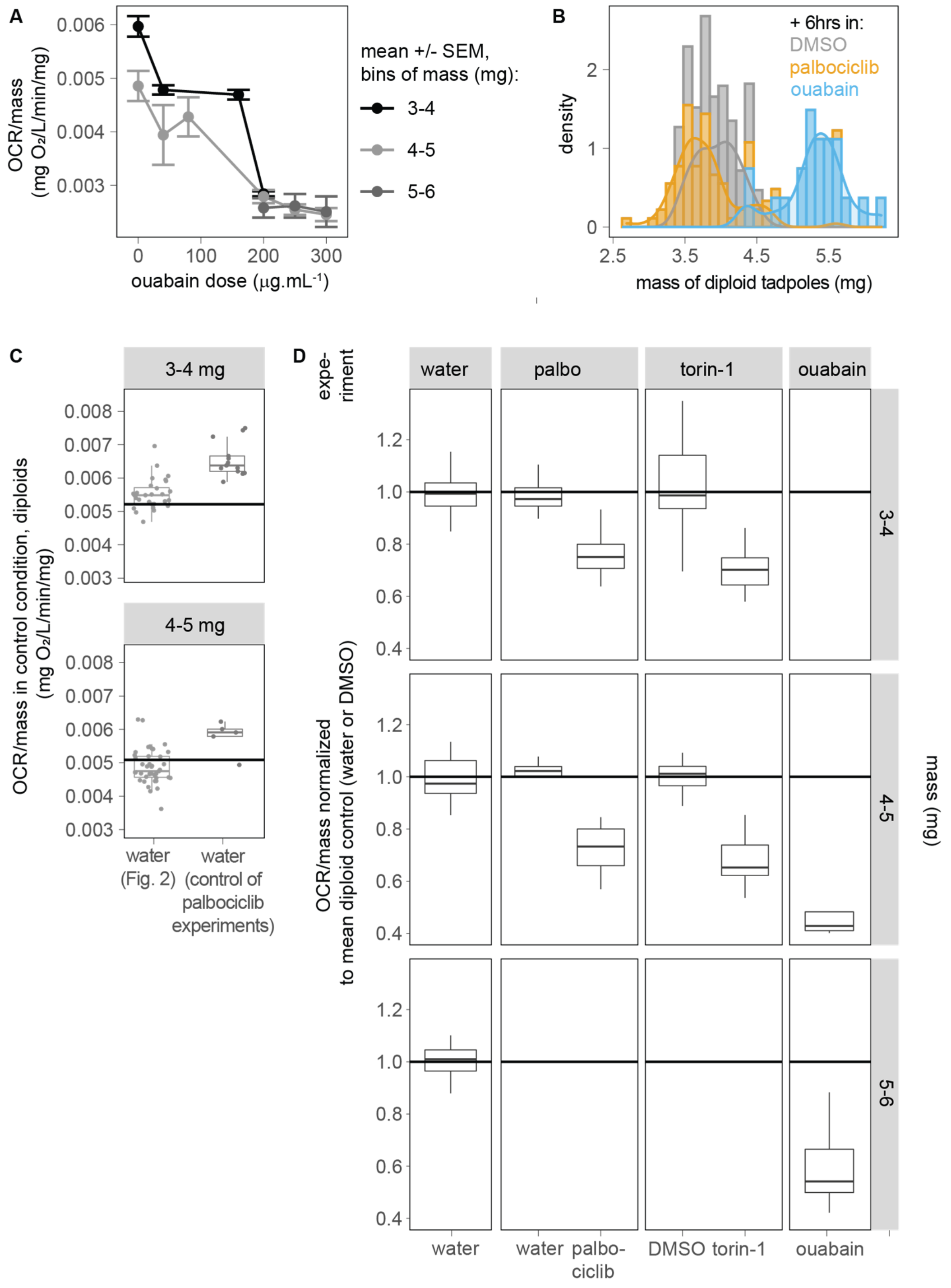


**Fig. S5. Details of ouabain experiments and normalization approach for estimation of the fraction of growth and maintenance to total embryo energy. (A)** OCR per mass as a function of ouabain doses for stage 41 diploid embryos and two bins of mass. **(B)** The 6-hr treatment with ouabain induced a significant increase in embryo body mass compared with embryos treated with palbociclib or DMSO. This is likely because of perturbations of intracellular osmolarity following inhibition of the $Na^+/K^+$ ATPase. **(C)** The average OCR measured in diploids was a higher in untreated controls (in water) of the set of experiments performed for the palbociclib assay (Fig.

3C) compared with previous measurements in water (Fig. 2B-C). This may be due to a transient problem in calibration of the SDR device since later experiments in water or DMSO (Fig. 3D) in diploids were again at the initially measured value. Hence, to calculate the fraction of OCR decrease as a function of drug treatments in Fig. 3E, we first normalized each drug-treated condition to the mean diploid untreated value for that experiment **(D)**.

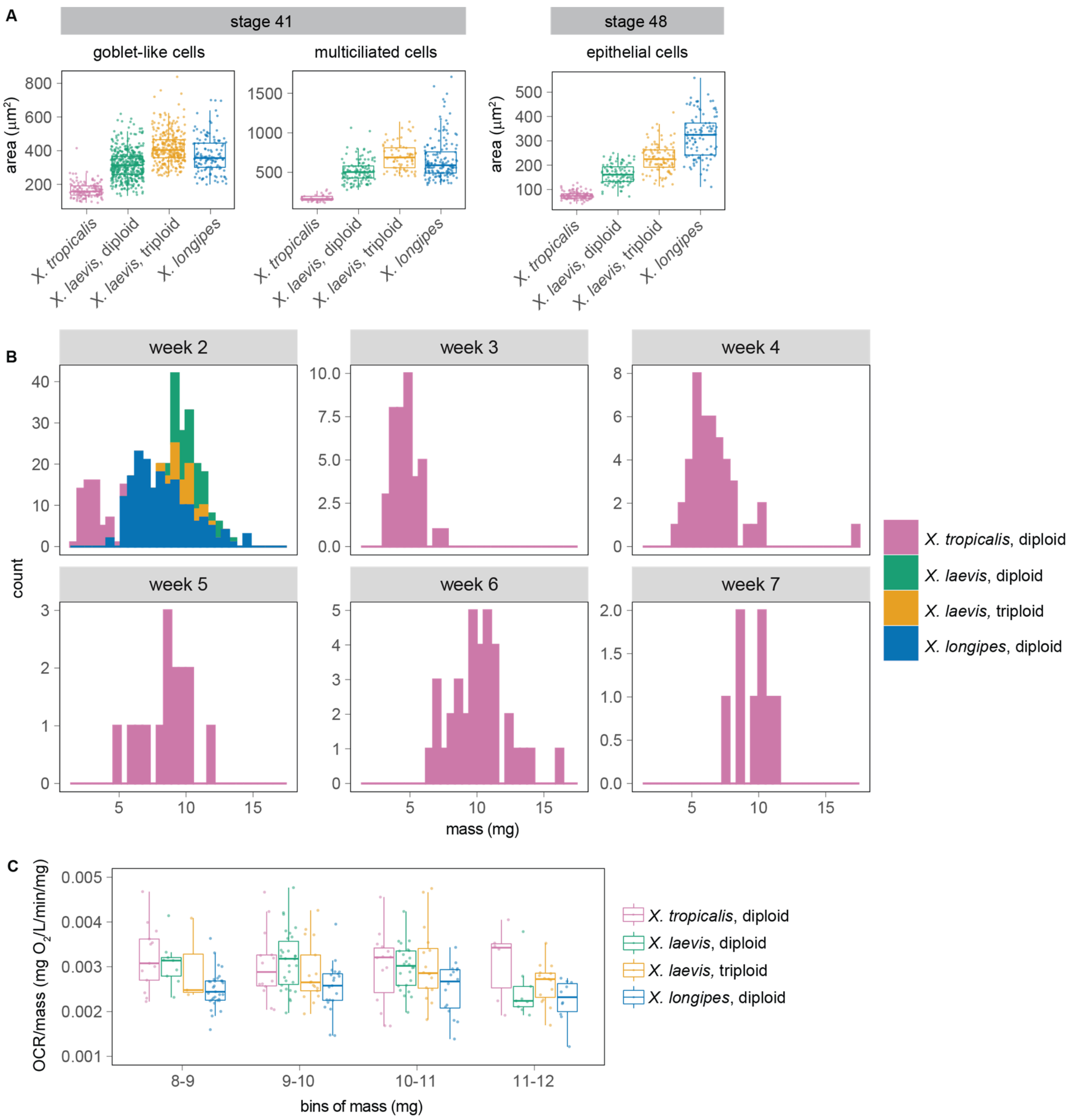


**Fig. S7. Details of cell size and mass measurements across species.** (**A**) Area of goblet-like and multicilliated cells measured as in Fig. 2A-B at stage 41 and at stage 47 (data for *X. tropicalis* and *X. longipes* at stage 47 come from ref.(*37*)). **(B)** Distribution of tadpole mass across species shows that *X. laevis* and *X. longipes* have overlapping masses during week 2 while *X. tropicalis* did not. In Fig. 4C-D, to compare tadpoles of similar mass, *X. tropicalis* from week 5-7 are compared with *X. laevis* and *X. longipes* from week 2 (see also Table S5). **(C)** Same as in Fig. 4C but with the largest bin of mass, 11-12 mg. *X. laevis* diploids displayed a lower OCR at this mass for unexplained reasons, but otherwise trends among *X. tropicalis,* triploid *X. laevis* and *X. longipes* appeared qualitatively similar to that observed for all other mass bins.
